# BRG1 establishes the neuroectodermal chromatin landscape to restrict dorsal cell fates

**DOI:** 10.1101/2023.06.24.546365

**Authors:** Jackson A Hoffman, Ginger W Muse, Lee F Langer, Isabella Gandara, James M Ward, Trevor K Archer

**Author notes:** These authors contributed equally (Jackson A Hoffman, Ginger W Muse).

## Abstract

Cell fate decisions are achieved with gene expression changes driven by lineage-specific transcription factors (TFs). These TFs depend on chromatin remodelers including the BAF complex to activate target genes. BAF complex subunits are essential for development and frequently mutated in cancer. Thus, interrogating how BAF complexes contribute to cell fate decisions is critical for human health. We examined the requirement for the catalytic BAF subunit BRG1 in neural progenitor cell (NPC) specification from human embryonic stem cells. During the earliest stages of differentiation, BRG1 was required to establish chromatin accessibility at neuroectoderm-specific enhancers. BRG1 depletion resulted in abnormal NPC populations that differentially expressed neuroectodermal TFs, were more prone to neuronal differentiation, and precociously formed neural crest lineages. These findings demonstrate that BRG1 mediates NPC specification by ensuring proper expression of lineage-specific TFs and appropriate activation of their transcriptional programs.

## INTRODUCTION

Access to DNA is regulated by large multi-subunit chromatin remodeling complexes that work in concert with many nuclear processes to ultimately regulate gene expression and cell identity. The SWItch/Sucrose Non-Fermentable (SWI/SNF, also known as BAF) complex utilizes ATP hydrolysis to slide or eject histone octamers to remodel nucleosomes^1^. In general, the BAF complex is required to maintain chromatin accessibility at promoters and regulatory elements to facilitate transcription factor and transcriptional machinery interactions^2, 3^. As such, the BAF complex is associated with and often required for active transcription.

Understanding how BAF complexes regulate cell identity and cell fate transitions is of great interest to human health research. It is estimated that >20% of human cancers contain mutations in at least one BAF subunit^4, 5^, and loss of BRG1 protein expression is reported to occur in >40% of brain, liver, kidney and intestinal cancers^6^. The BAF complex is also essential throughout development, and loss of BRG1 (or most other BAF subunits) results in embryonic lethality ^7^. Conversely, mutation or dysregulation of BAF subunits can also have oncogenic effects, and high levels of BRG1 expression are found in a variety of tumor types and often associated with poorer prognoses^8–10^. Thus, the activity and expression of the BAF complex is essential and there are context-dependent requirements for BAF subunits throughout development and in tumorigenesis.

Mutations in BRG1 or the core subunit BAF47 can give rise to highly aggressive pediatric brain tumors such as atypical teratoid/rhabdoid tumors (ATRTs) and medulloblastomas ^11–17^. In the case of ATRTs, the requirement for BAF47 is strongly established and BAF47 mutations are found in the vast majority of tumors^18^. BRG1 mutations in ATRT are rarer and associated with earlier onset and worse prognoses^19–21^. As these tumors are thought to arise from progenitor cells during nervous system development, a comprehensive understanding of BAF complex functions during neuroectodermal differentiation is critical and human embryonic stem cells (ESCs) provide an apt model system for investigation.

Individual BAF subunits have a variety of context-dependent requirements during ESC self-renewal and differentiation in both human and mouse^22^. Previously, we found that BAF47 is essential for neuroectodermal differentiation of human ESCs, during which it is required to regulate enhancer accessibility^23^. Knockdown of either BRG1 or BAF47 in ESC leads to de-repression of bivalent lineage-specific genes, suggesting that the BAF complex safeguards the pluripotent state of self-renewing ESC^23, 24^. During neuroectodermal differentiation, the BAF complex undergoes a series of tightly regulated paralogue switching events, with distinct subunit compositions defining ESC-, neural progenitor-, and neuron-specific complexes^25^. Mutation of the complex-specific subunits results in corresponding defects in pluripotency, differentiation, and neural morphogenesis and function^26–29^. In general, conditional deletion of BAF subunits during specific stages of neural differentiation leads to defective proliferation and specification of neural cell types^30^. Throughout these processes, the BAF complex can interact with a variety of lineage-determining neural transcription factors such as SOX2, PAX6, and ASCL1^31–34^.

We set out to directly assess the requirement for the BRG1 subunit and BAF complex activity in self-renewing ESC and during neural progenitor cell (NPC) specification. By depleting BRG1 with either shRNA expression or PROTAC (proteolysis targeting chimera) treatment, we demonstrate that BAF complex activity is essential for proper patterning of NPCs at the onset of neuroectodermal differentiation. BRG1 depletion resulted in the dorsalization of NPCs, enhanced and altered neuronal differentiation, and precocious neural crest differentiation. BRG1 was required to establish and maintain accessibility at enhancers that were activated during the initial stages of differentiation. These BRG1-dependent enhancers were highly enriched for neuroectodermal lineage transcription factor motifs. Thus, we concluded that BAF complex activity is essential for the establishment of the NPC chromatin landscape by lineage-specific transcription factors.

## RESULTS

### BRG1 controls neurectoderm developmental transcription programs in ESC

Inducible shRNA depletion of BAF47 protein in H1 human embryonic stem cells (ESC) resulted in widespread de-repression of gene expression and inhibition of directed neuroectodermal differentiation^23^. This demonstrated a critical role for the BAF47-containing BAF complexes in ESC and during differentiation. However, BRG1 immunoprecipitation followed by mass spectrometry revealed that BAF complex assembly was not disrupted by the reduction of BAF47 protein (Figure S1). Independent of BAF47 depletion, BRG1 specifically pulled-down BAF subunits from all three variants of the BAF complex (BAF, PBAF, and ncBAF, Figure S1). Thus, it was unclear whether the changes observed in BAF47KD ESC were the result of altered, reduced, or mis-targeted BAF complex activity.

To directly investigate the requirement for BAF complex activity in ESC and neuroectodermal differentiation, we generated an H1-derived cell line with inducible expression of BRG1 shRNA (BRG1KD cells). Doxycycline induction of BRG1 shRNA expression for 72 hours resulted in 80- 90% depletion of BRG1 protein levels with minimal impact on the expression of pluripotency factors (Figure 1A, S2). BRG1 depletion resulted in 1,477 differentially expressed genes (DEGs, fold change >1.5, adjusted p-value <0.05) (Figure 1B, S2). Two-thirds of these DEGs were down- regulated after BRG1 depletion, consistent with the association of BRG1 activity and transcriptional activation. Despite this bias towards a requirement for transcriptional activation, 499 genes were up-regulated following BRG1 depletion. BRG1 depletion also significantly altered the transcriptional at 498 ESC enhancers^35^, with nearly all of these being down-regulated (Figure 1C). Thus, BRG1 was predominantly required to promote gene expression in ESC.

**Figure 1:**
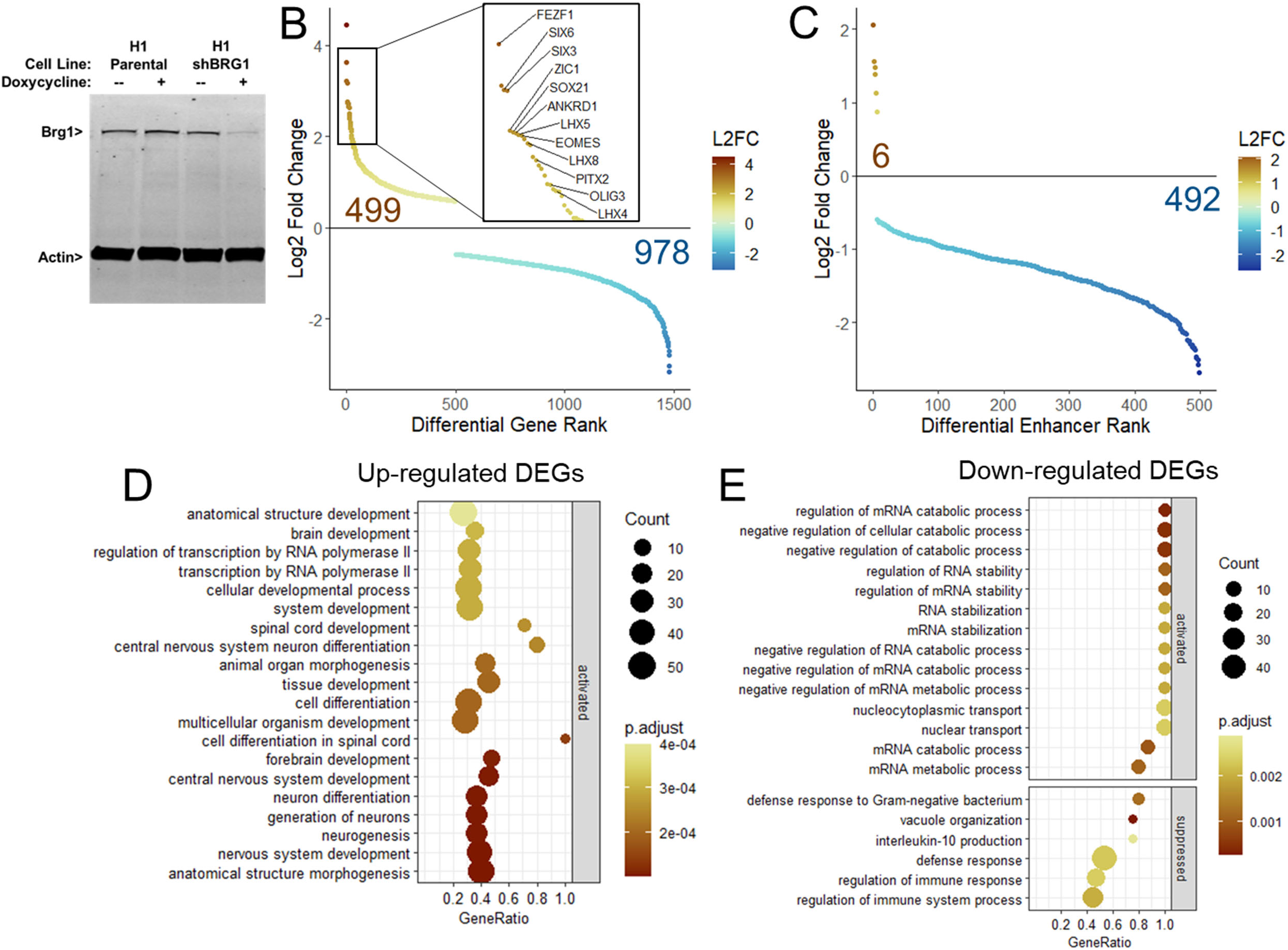
BRG1 controls neurectoderm developmental transcription programs in ESC. A) Western blot for BRG1 and Actin protein levels in parental and BRG1KD cells +/- Dox. B) Mean RNAseq log2 fold change of ranked DEGs following BRG1KD. Inset highlights select up- regulated neuroectodermal transcription factors. See also Figure S2. C) Mean RNAseq log2 fold change of ranked enhancers following BRG1KD. D) Gene set enrichment plot of basic processes enriched in up-regulated BRG1KD DEGs. E) Gene set enrichment plot of basic processes enriched in down-regulated BRG1KD DEGs.

Previously we showed that transcription factors involved in neural development such as ZIC1, SOX21, and FEZF1 were among the most highly upregulated genes in BAF47KD ESC^23^. These genes were among 12 transcription factors relevant to neuroectodermal differentiation within the top 25 genes up-regulated in BRG1KD ESC (Figure 1B inset, S2). Additionally, the up-regulated DEGs were significantly enriched for biological pathways corresponding to activation of neural differentiation and development (Figure 1D). Conversely, down-regulated DEGs were enriched for pathways involved in mRNA metabolism and immune response (Figure 1E). Thus, the transcriptomic changes observed in BRG1KD ESC suggested that BRG1 may also play a role in neuroectodermal differentiation.

### BRG1 depletion promotes formation of atypical neural progenitor cell populations

To determine the effect of BRG1 depletion on neuroectoderm differentiation, we utilized dual SMAD inhibition^36^ to induce the formation of neural progenitor cells (NPC) over the course of a nine-day differentiation protocol (Figure 2A). To ensure depletion of BRG1 protein levels at the onset of differentiation, BRG1KD ESC were pre-treated with Doxycycline (Dox) for 72 hours and then split into neural induction medium (NPC media). Dox was maintained in NPC media throughout the protocol, ensuring ongoing depletion of BRG1 protein (Figure 2B). Both control and BRG1KD cells exhibited near complete loss of OCT4 and NANOG expression by day 3 in NPC media, indicating that BRG1KD cells were able to successfully exit pluripotency and begin differentiating (Figure S3B, D). In contrast, we previously showed that despite similar upregulation of neuroectodermal transcription factors, BAF47KD ESC were resistant to neural induction and maintained ESC characteristics^23^. Thus, BRG1 knockdown was distinct from BAF47 knockdown during the first stages of NPC differentiation.

**Figure 2:**
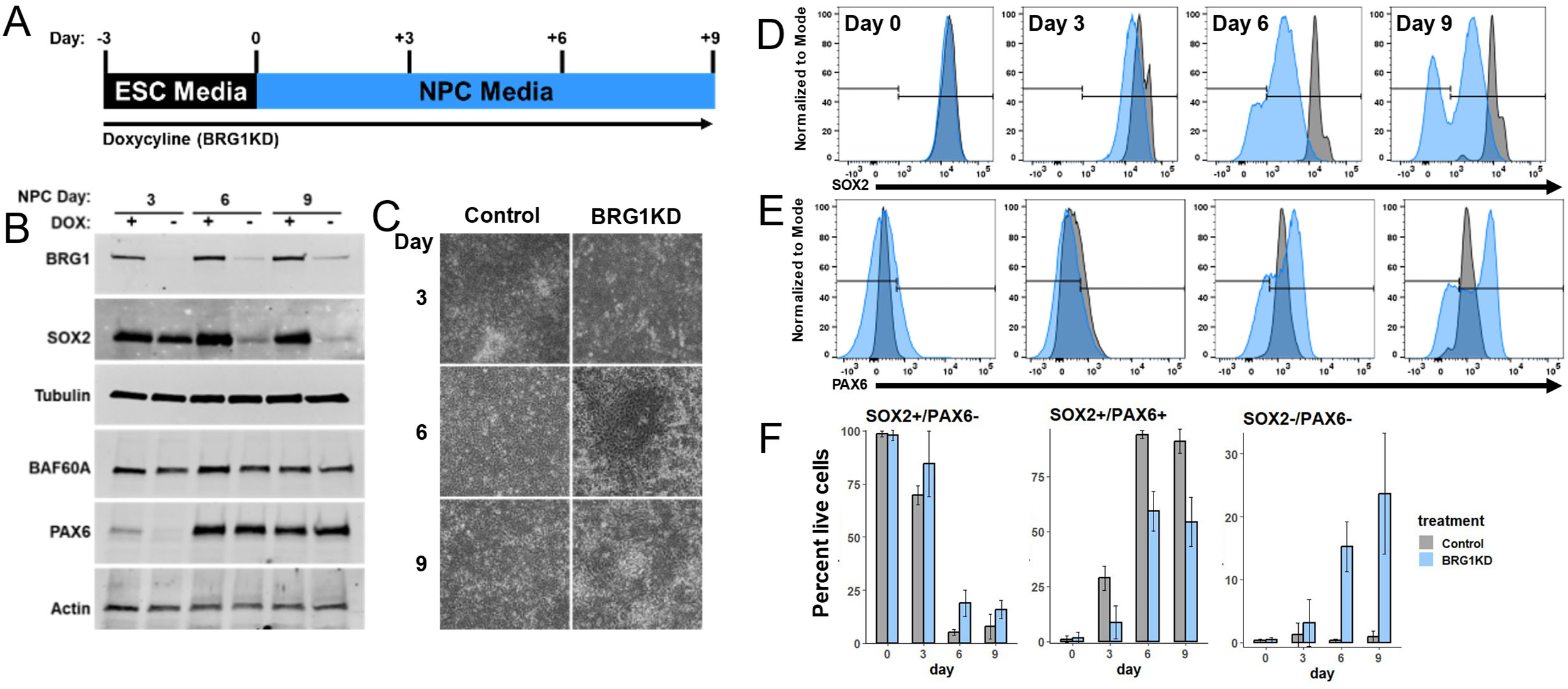
BRG1 depletion promotes formation of atypical neural progenitor cell populations. A) Graphical depiction of experimental setup for panels B-F. B) Western blots showing BRG1, SOX2, Tubulin, BAF60A, PAX6, and Actin protein expression at collection timepoints. See also Figures S2 and S3. C) Phase contrast images of control and BRG1KD NPCs. D-E) FACS histogram of SOX2 (D) and PAX6 (E) protein detection at collection timepoints. Log10 fluorescent signal depicted for control cells in gray and BRG1KD cells in light blue. Data represents the total of all biological replicates (n >= 3). F) Graphs depicting the percent of cells at each collection timepoint that were SOX2+/PAX6-, SOX2+/PAX6+, or SOX2-/PAX6-. Bar height indicates mean percentages and error bars represent standard deviation of biological replicates (n >= 3). Control values in gray, BRG1KD values in light blue.

Over the course of the NPC differentiation protocol, control cells formed a dense lawn of cells (Figure 2C). By day 6 of differentiation, BRG1KD cultures had a more heterogeneous appearance with frequent regions of cells with mesenchymal morphology (Figure 2C). This abnormal morphology suggested that BRG1 depletion had altered the normal course of differentiation. NPC differentiation can be tracked by measuring the expression of SOX2 and PAX6, two transcription factors that are critical for NPC specification and maintenance. In control cells, SOX2 was highly expressed in ESC and maintained at high levels through day 9 of NPC differentiation (Figure 2D). PAX6 was not expressed in control ESC and became uniformly expressed by day 6 of NPC differentiation (Figure 2E). By days 6 and 9, nearly all control cells were positive for both SOX2 and PAX6 (Figure 2F, S3C). BRG1KD cells also formed populations of PAX6 or SOX2 positive NPCs, however, BRG1KD atypically resulted in the presence of cells lacking the expression of these two markers. BRG1KD ESC expressed high levels of SOX2 prior to differentiation, but levels of SOX2 expression were progressively diminished over the course of NPC differentiation (Figure 2D). By day 9, two distinct populations of cells formed in BRG1KD cells – a SOX2+ population that maintained SOX2 expression (albeit at a lower level than control cells), and a SOX2- population with a complete absence of SOX2 expression (Figure 2D). SOX2 expression was also markedly reduced in BRG1KD NPCs when detected via western blot or RTPCR (Figure 2B, S3C). Similarly, PAX6 was not expressed in BRG1KD ESC and was upregulated during NPC differentiation (Figure 2E). However, instead of a uniformly PAX6+ cell population, BRG1KD cells exhibited a broader distribution of PAX6 levels, with a subset of cells expressing higher levels of PAX6 than observed in control cells (Figure 2E). At days 6 and 9, the majority of BRG1KD cells formed a similar SOX2+ PAX6+ population with lower levels of SOX2 expression and higher levels of PAX6 (Figure 2F, S3C). However, more than a third of BRG1KD cells instead formed SOX2+/PAX6- or SOX2-/PAX6- cell populations (Figure 2F, S3C). Thus, BRG1KD ESC formed atypical populations of cells during directed NPC differentiation.

### Dorsalization of BRG1KD NPCs and precocious neural crest differentiation

To interrogate the identity of the abnormal NPC populations formed from BRG1KD ESC, we next performed single-cell RNAseq (scRNAseq) with two biological replicates at days 3, 6, and 9 of NPC differentiation. A high level of consistency was observed between replicates at each timepoint, so replicates were merged and treated as individual timepoints for subsequent analyses. Following uniform manifold approximation and projection (UMAP) for dimensional reduction, BRG1KD NPCs formed clusters that were almost entirely distinct from control NPCs (Figure 3A). Control NPCs from each timepoint formed defined and separate clusters (Figure 3B). Conversely, day 3 BRG1KD NPCs formed a distinct cluster but day 6 and 9 BRG1KD NPCs formed a single overlapping cluster (Figure 3B). Similar to our FACS analyses, control NPCs were uniformly positive for both PAX6 and SOX6 (Figure 3C,D). At day 3, BRG1KD NPCs expressed lower levels of SOX2 and lacked upregulation of PAX6 (Figure 3C,D). By days 6 and 9, multiple populations arose in BRG1KD NPCs: 1) SOX2+/PAX6-high cells, 2) SOX2+/PAX6- cells, and 3) SOX2-/PAX6- cells (Figure 3C,D). Thus, scRNAseq enabled the identification of the distinct cell populations observed among BRGKD NPCs.

**Figure 3:**
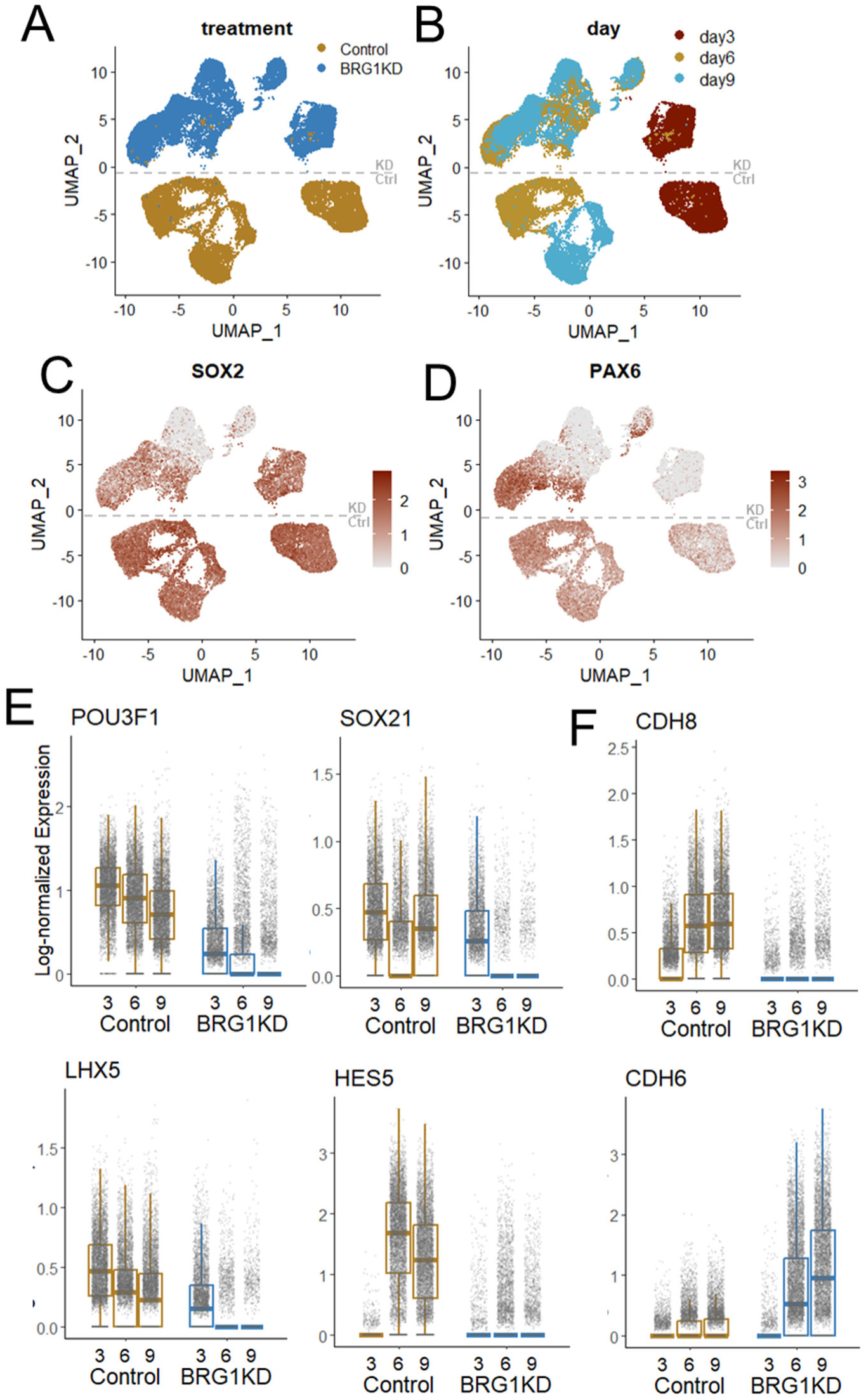
Single cell RNA-seq captures abnormal cell populations in BRG1KD NPCs. A-B) UMAP plots depicting clustering of NPCs by treatment (A) and day (B). C-D) UMAP feature plots depicting SOX2 (C) and PAX6 (D) log-scaled expression. E-F) Box-and-jitter plots depicting log-scaled expression of neuroectodermal transcription factors (G) and cadherins (H) in control and BRG1KD at days 3, 6, and 9. Each dot represents a single cell, box edges represent the 25^th^ and 75^th^ percentiles and midline represents the median.

Examination of the expression of additional NPC marker genes further revealed the abnormality of BRG1KD NPCs. Most control NPCs expressed transcription factors involved in NPC maintenance and identity such as POU3F1, SOX21, LHX5, and HES5 (Figure 3E, 4C). Conversely, expression of these markers was absent from the majority of BRG1KD NPCs (Figure 3E). Similarly, control NPCs expressed CDH8, a cell surface marker of neurogenic progenitors in the developing brain^37^ (Figure 3F). Most BRG1KD NPCs lacked CDH8 expression and instead expressed CDH6, a marker of gliogenic progenitors (Figure 3F)^37^. As such, BRG1KD NPCs appeared to be diverging from the typical NPC phenotype towards an alternative cell fate.

To gain further insight into the potential fates of BRG1KD NPCs, we performed cell type annotation using CellTypist^38^ and an atlas of transcriptional profiles from first-trimester fetal human brains^39^. Most control NPCs were annotated as telencephalon or forebrain radial glia, demonstrating that control NPCs adopted a ventral or rostral cell fate (Figure 4A). At day 3, BRG1KD NPCs were also annotated as telencephalon radial glia, indicating that BRG1KD NPCs began differentiation on the same fate trajectory as control NPCs (Figure 4A). However, at days 6 and 9, BRG1KD NPCs were mostly annotated as midbrain, hindbrain, and medulla radial glia, indicating that BRG1 depletion had resulted in NPCs with a more dorsal or caudal NPC fate. Indeed, BRG1KD NPCs had strongly up-regulated expression of transcription factors associated with the development of the dorsal neural tube such as ZIC1, PAX3, and MSX1 (Figure 4B,C, S3D). Surprisingly, a subset of BRG1KD NPCs were also annotated as “head neural crest cells” and “brain placode cells” (Figure 4A). This further indicated that BRG1KD NPCs had adopted a fate similar to that of cells at the dorsal side of the developing neural tube from which the neural crest arises or the dorsal ectoderm that gives rise to the placodes.

**Figure 4:**
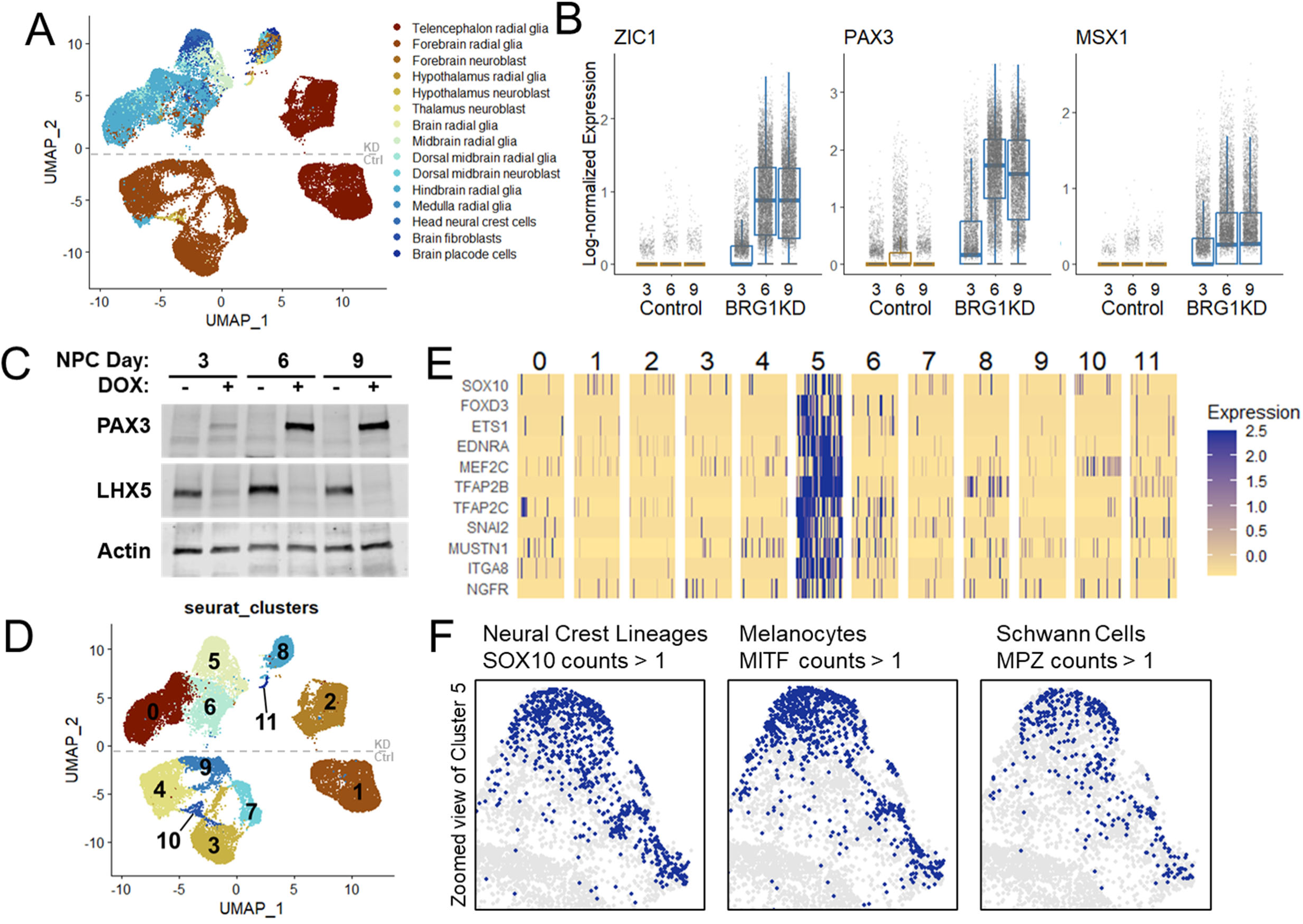
Dorsalization of BRG1KD NPCs and precocious neural crest differentiation. A) UMAP plot depicting first-trimester fetal human brain-based cell type annotations. B) Box-and- jitter plots of log-scaled expression of dorsal neuroectodermal transcription factors. at days 3, 6, and 9. Each dot represents a single cell, box edges represent the 25^th^ and 75^th^ percentiles and midline represents the median. C) Western blots for PAX3 and LHX5 at NPC collection timepoints. D) UMAP plot showing de novo Seurat cluster assignments. See also Figures S4 and S5. E) Heatmap of log-scaled expression of neural crest cell marker genes in randomly down-sampled Seurat clusters (50 cells per cluster). F) Zoomed-in UMAP plots highlighting cells in cluster 5 with more than 1 count of SOX10, MITF, and MPZ.

To facilitate the characterization of both control and BRG1KD cell populations, we used de novo clustering to define 11 cell clusters (Figure 4D). Among BRG1KD-specific clusters, cluster 5 included the BRG1KD cells annotated as neural crest. To confirm that these BRG1KD cells had adopted a neural crest fate, we examined marker genes that were specifically enriched in cluster 5. Indeed, markers of neural crest cells such as FOXD3, TFAP2B, and NGFR were strongly enriched in cluster 5 (Figure 4E). A subset of cluster 5 cells were also enriched for SOX10, a marker of further neural crest-derived lineages (Figure 4E). Strikingly, some BRG1KD cells in cluster 5 also expressed MITF and MPZ, marker genes of melanocytes and Schwann cells, respectively (Figure 4F). Thus, BRG1 depletion promoted atypical differentiation of NPCs to neural crest cells and more specialized neural crest-derived cell types.

Clusters 8 (BRG1KD cells) and 10 (control cells) were readily identifiable as clusters that exhibited neuronal gene expression patterns (Figure S4A,C). Both clusters were specifically enriched for a variety of neuronal lineage markers such as NHLH1, STMN2, and ELAVL3 (Figure S4C). Cluster 10 accounted for 1.7% and 5.1% of control cells at days 6 and 9, respectively (Figure S4B). BRG1KD NPCs appeared to be more prone toward neuronal differentiation, as cluster 8 was comprised of ∼10% of BRG1KD cells at days 6 and 9 (Figure S4B). However, the BRG1KD cluster also exhibited differential neuronal gene expression from the control cluster, with reduced/absent expression of genes such as PRMT8 and NEUROD4 and increased expression of ASCL1, SNCG, and GYG1 (Figure S4D,E). This demonstrated that BRG1KD NPCs more readily progressed to neuronal fates and formed neuronal cells with atypical gene expression patterns.

Cluster 11 also represented a unique BRG1KD cell cluster with a more advanced cell fate than expected. Comprised of 88 day 9 and 2 day 6 cells, this cluster was uniquely enriched for markers of cilia development and morphogenesis such as CCNO, CDC20B, RSPH1, and several genes in the CFAP family (Figure S5A, B). Gene set enrichment analysis confirmed that genes involved in cilium assembly were strongly enriched among cluster 11 marker genes (Figure S5C). Thus, depletion of BRG1 during NPC differentiation permitted ectopic specification of NPC-derived ciliated cells, potentially representative of polarized neuroepithelial cells or ciliated neurons^40^. Taken together, cell annotation and cluster identification revealed that BRG1KD NPCs adopted a dorsalized cell fate that uniquely gave rise to neural crest cells and was more prone to neuronal differentiation.

### BRG1 is required during the initial stages of neural progenitor cell specification

We next sought to validate and refine the requirement for BRG1 during early NPC differentiation. We utilized ACBI1, a BRG1/BRM/BAF180 proteolysis targeting chimeric (PROTAC) protein degrader, for acute depletion of BRG1^41^. In ESC, BRG1 protein was rapidly degraded by PROTAC treatment (Figure 5A). We next repeated our NPC differentiation and FACS experiments +/- 24 hours of PROTAC. PROTAC was maintained in the culture media for the duration of the differentiation time course (Figure 5B). PROTAC treatment largely recapitulated the effects of BRG1KD and resulted in non-uniform expression of PAX6 and SOX2 in day 6 NPCs (Figure 5C, S6A,B). Like BRG1KD NPCs, PROTAC NPCs formed a substantial population of SOX2-/PAX6- cells (Figure 5C, S6B). However, PROTAC NPCs did not form a distinct SOX2+/PAX6- population as observed in BRG1KD NPCs (Figure S6B). PROTAC treatment also reproduced the upregulation of the dorsal transcription factors PAX3 and MSX1 observed in BRG1KD NPCs (Figure 5D). To validate the formation of neural crest cells in BRG1-depleted NPCs, we utilized the neural crest marker NGFR. In BRG1KD NPCs, 9.3% of cells were NGFR+ by flow cytometry, and these NGFR+ cells were also SOX2-/PAX6- (Figure 5C, S6C,E). PROTAC treatment yielded similar results, with NGFR+ cells comprising 8.8% of the total number of cells (Figure 5D, S6D). Taken together, these experiments demonstrated that the BRG1KD cell populations identified by scRNAseq could be validated by FACS and largely recapitulated by PROTAC-mediated depletion.

**Figure 5:**
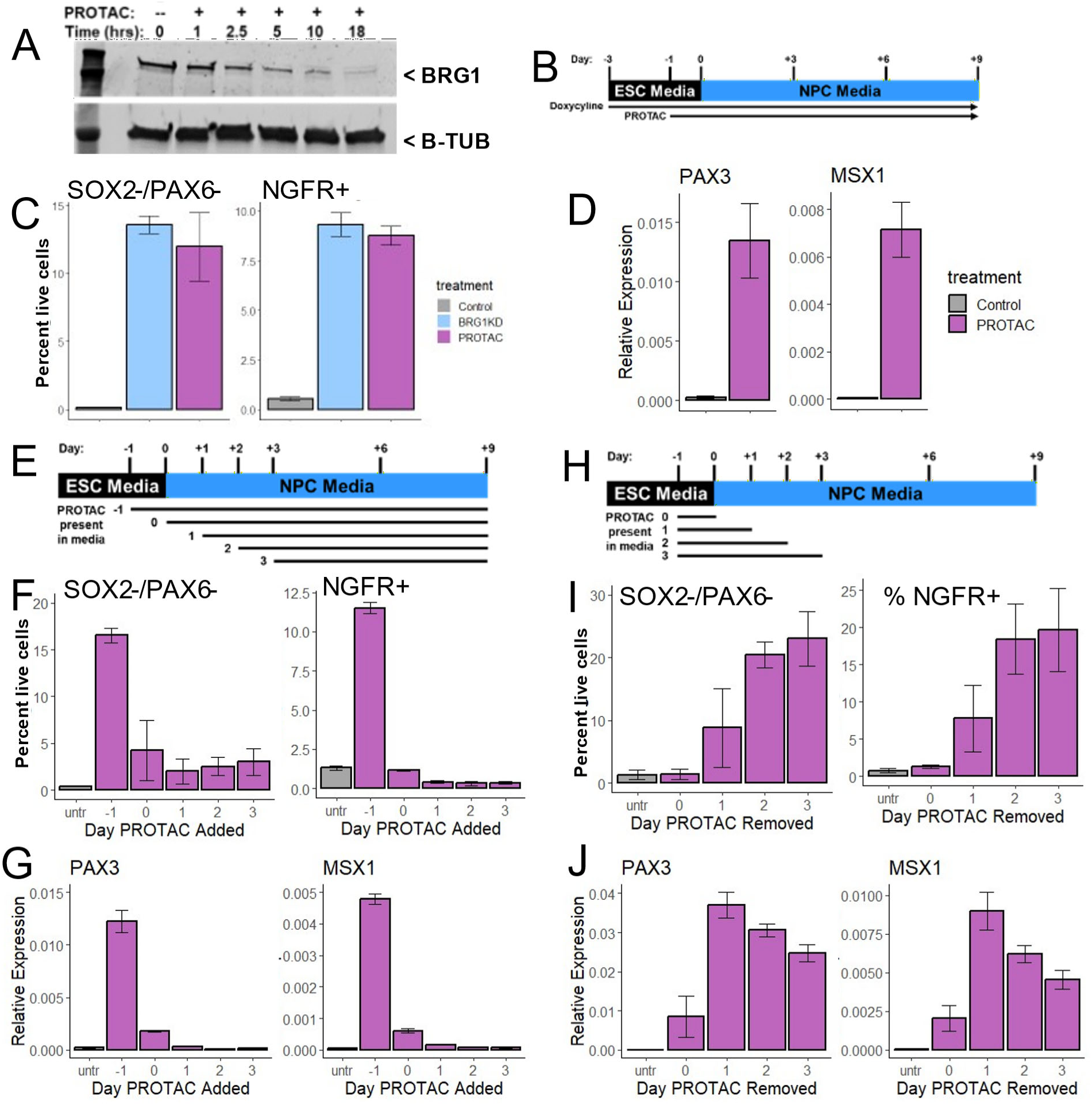
BRG1 is required during the initial stages of neural progenitor cell specification. A) Western blot of BRG1 and beta-tubulin protein levels over a time course of PROTAC treatment. B) Graphical depiction of experimental setup for C,D. See also Figure S6. C) Percent of cells in each treatment that were detected as SOX2-/PAX6- and NGFR+ by FACS at day 6. D) RT-PCR for PAX3 and MSX1 at day 6. E) Graphical depiction of experimental setup for F-G. See also Figure S6. F) Percent of cells in each treatment that were detected as SOX2-/PAX6- and NGFR+ by FACS at day 9. G) RT-PCR for PAX3 and MSX1 at day 9. H) Graphical depiction of experimental setup for I,J. See also Figure S6. I) Percent of cells in each treatment that were detected as SOX2-/PAX6- and NGFR+ by FACS at day 9. J) RT-PCR for PAX3 and MSX1 at day 9. For C,F, and I, bar heights indicate mean percentages and error bars represent standard deviation of biological replicates (n >= 3). For D, G, and J, bars depict mean expression level relative to geometric mean of ACTB, GAPDH, and TUBA1B; error bars represent standard deviation of biological triplicates.

As PROTAC treatment provided for more acute depletion of BRG protein, we designed experiments to identify the window during which BRG1 was required for typical dual-SMAD inhibition NPC differentiation. We first designed an experiment to determine whether BRG1 was required at the onset of NPC specification (Figure 5E). For our standard experimental conditions PROTAC treatment was initiated 1 day prior to the switch to differentiation media. We defined this as addition at day -1 and further defined a series of treatment schemes with the day of PROTAC treatment initiation ranging from day 0 to day 3 (Figure 5E). Using day 9 as the endpoint, only the standard “T=-1” scheme yielded a robust population of SOX2-/PAX6- cells (Figure 5F, S6F,H). Beginning PROTAC treatment at later timepoints resulted in a diminished SOX2-/PAX6- population, with at most a third of the cells detected with the -1 timepoint (Fig5F, S6H). Similarly, NGFR+ neural crest cells and upregulation of PAX3 and MSX1 were robustly detected when PROTAC was added at day -1, but not detected above control levels in the other treatment schemes (Figure 5F,G, S6H,I). We thus concluded that the SOX2-/PAX6- and NGFR+ cell populations required BRG1 depletion at the onset of NPC differentiation.

We next examined the duration of PROTAC treatment required to produce the BRG1 depletion phenotype. To do so, we control and constant PROTAC treatment NPCs to samples in which PROTAC was added at day -1 for all samples, but then removed at day 0, 1, 2, or 3 (ie after 24, 48, 72, or 96 hours of treatment, Figure 5H). As with constant treatment, removal of PROTAC at day 2 or day 3 resulted in robust populations of SOX2-/PAX6- cells and NGFR+ neural crest cells (Figure 5I, S6J,K). The size of these cell populations were reduced by more than half when PROTAC was removed at day 1, and completely lost when PROTAC was removed at day 0 (Figure 5I, S6J,K). Robust upregulation of PAX3 and MSX1 occurred a day earlier in the time course, with partial induction in the day 0 timepoint and full induction with the day 1 timepoint (Figure 5J). Thus, PROTAC treatment needed to be maintained for at least 48 hours to induce full upregulation of dorsal markers and for 72 hours to recapitulate the SOX2-/PAX6- and NGFR+ cell populations observed with constant treatment. Taken together, these experiments indicated that BRG1 depletion during the first 48-72 hours of NPC differentiation resulted in the formation of a dorsalized SOX2-/PAX6- NPC population with the capacity to form neural crest cell lineages.

### BRG1 maintains chromatin accessibility and enhancer marks at neuroectoderm enhancers

The BAF complex has typically been associated with open and active regions of chromatin. We thus hypothesized that BRG1 was required to either maintain or establish accessible chromatin at enhancers and/or promoters during the initial stages of NPC differentiation. To test this, we pre-treated cells with either 72 hours of Dox to induce BRG1KD or 24 hours of PROTAC to degrade BRG1 protein and maintained these treatments through the first 3 days of NPC differentiation (Figure 6A). We collected biological triplicates at day 0 and day 3 and performed ATAC-seq to identify accessible chromatin regions in ESC and day 3 NPCs. For analysis of accessible chromatin, nucleosome free regions (NFR) were defined as fragments <100 base pairs and NFR reads were used to call peaks from each replicate. BRG1KD and PROTAC samples consistently had fewer NFR peaks than control samples in both cell types indicating that BRG1 depletion resulted in a loss of accessible chromatin (Figure 6B).

**Figure 6:**
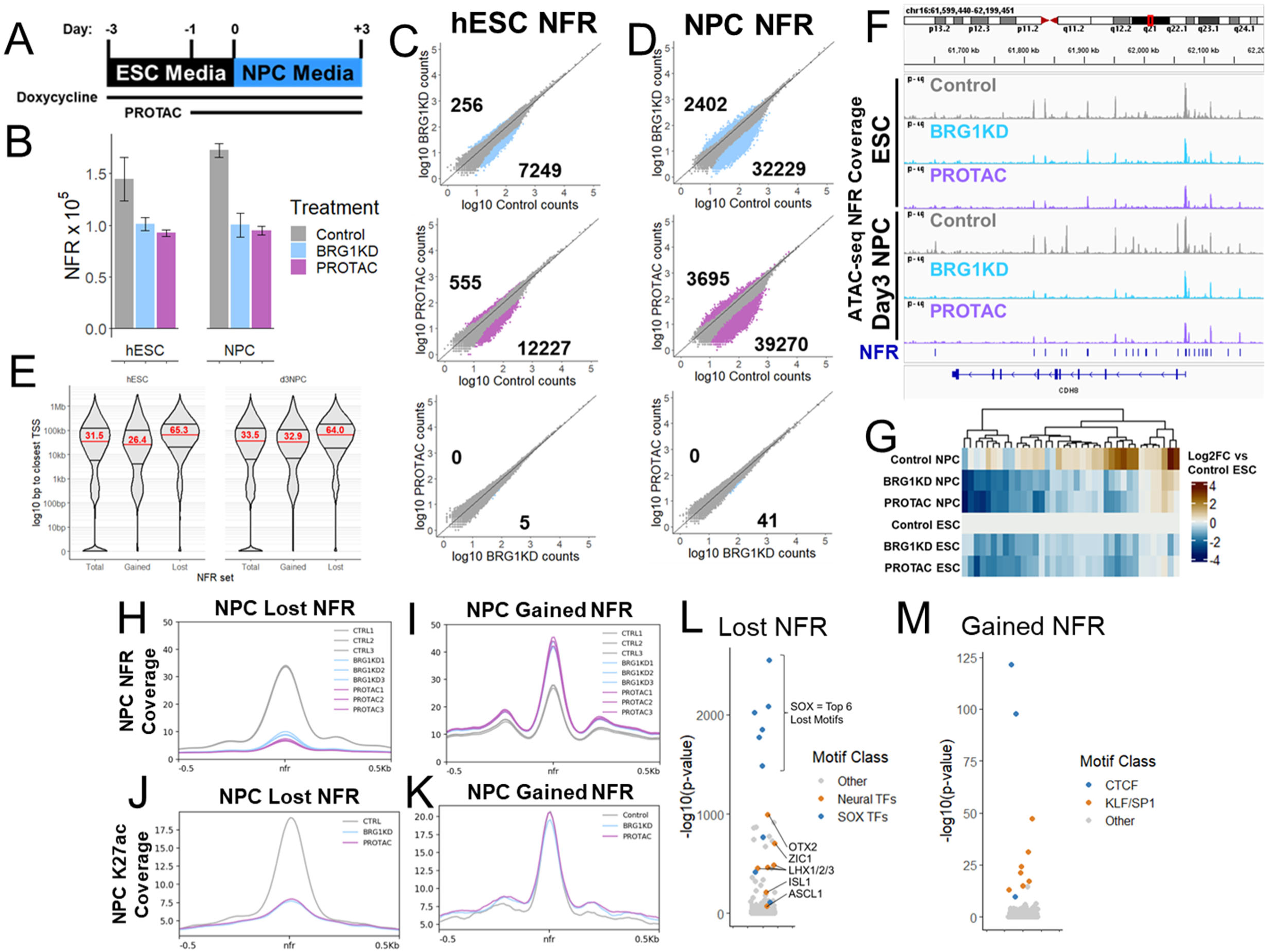
BRG1 maintains chromatin accessibility and enhancer marks at neuroectoderm enhancers. A) Graphical depiction of ATAC-seq experimental set-up. Cells were collected at day 0 (ESC) and day 3 (NPCs). B) Number of NFR called in each treatment/timepoint. Bars represent mean and error bars represent standard deviation of biological triplicates. C-D) Scatter plots comparing NFR counts between treatments over the total set of NFR in ESC (C) and NPCs (D). Colored dots and numbers represent NFR with significantly more or less accessibility. E) Violin plots of NFR distance from the closest annotated TSS for the total NFR set, NFR that gain accessibility upon BRG1-depletion (Gained), and NFR that lose accessibility upon BRG1-depletion (Lost). Red numbers represent the median distance for each NFR set. F) IGV browser snapshot of ATAC-seq NFR coverage around the CDH6 gene. NFR peaks are annotated as blue lines above the gene model. Scale for all tracks is 0-44. G) Heatmap of log2 fold change relative to Control ESC of NFR coverage over the peaks depicted in F. H-I) Meta- profiles of NFR coverage over lost NFR (H) and gained NFR (I). J-K) Meta-profiles of H3K27ac Cut&Tag coverage over lost NFR (J) and gained NFR (K). L-M) Dot plots of transcription factor motifs enriched in lost NFR (L) and gained NFR (M).

For downstream analysis, we trimmed the total number of peaks down to a high confidence set of 137,958 ESC NFR peaks and 164,265 NPC NFR peaks. These peak sets were merged into combined peak set of 186,627 NFR peaks for differential analysis. Differentially accessible NFR were then defined as peaks with >2 fold change and an adjusted p-value <0.01 between sets of biological triplicates. BRG1 depletion predominantly resulted in loss of accessibility at both ESC and NPC NFR, with over tenfold more “lost” NFR than “gained” NFR (Figure 6C,D). Comparatively, there were very few differential NFR between BRG1KD and PROTAC conditions, indicating that the two distinct methods of BRG1 depletion had consistent effects on chromatin accessibility (Figure 6C,D). Intriguingly, the effect of BRG1 depletion was more pronounced in NPCs, with >3-fold more differential NFR in NPCs (Figure 6C,D). Thus, BRG1 was required to maintain accessible chromatin at NFR in both ESCs and NPCs.

To assess the genomic context of the differential NFR, we examined the distances between NFR and annotated TSSs. The genomic distribution of NFRs in ESC and NPCs was very similar. With the total sets of NFR at each timepoint, approximately 20% of peaks occurred within 1kb of an annotated TSS, the remaining 80% of peaks had a broad distribution ranging from >1kb to >1Mb from the nearest TSS, and the median distance to the closest TSS was ∼20kb (Figure 6E). In general, differential NFR did not directly overlap TSSs (Figure 6E). Despite this, the median distance to the closest TSS for NFR that gained accessibility following BRG1 depletion remained ∼20kb (Figure 6E). Conversely, NFR that lost accessibility after BRG1 depletion trended much further away from TSSs, with a median distance of ∼70kb (Figure 6E). For example, 37 NFR were called in a 600kb region surrounding the CDH8 gene, which is down-regulated in BRG1KD NPCs (Figure 6F, 3F). Accessibility was unaffected at the CDH8 TSS, but the accessibility of many of the surrounding NFR was significantly reduced following BRG1 depletion in both ESC and NPCs (Figure 6F,G). The extent of accessibility changes at gained and lost NFR also differed, with a greater magnitude of change in accessibility at lost NFR (Figure 6H,I). Both gained and lost NFR were marked by H3K27ac Cut&Tag enrichment in control cells, indicating that these NFR likely represented active enhancers (Figure 6J,K). However, at the lost NFR, BRG1 depletion resulted in near complete loss of H3K27ac enrichment, whereas enrichment was largely unchanged at gained NFR (Figure 6J,K). As such, BRG1 was required during the first three days of NPC differentiation to maintain accessibility and H3K27 acetylation at thousands of active enhancers.

To gain insight into the potential regulation of these BRG1-dependent enhancers, we searched for enriched transcription factor motifs within the NFR. The NFR that lost accessibility after BRG1 depletion in NPCs were most strongly enriched for SOX motifs (Figure 6L). The lost NFR were also enriched for motifs of other transcription factors involved in neuroectodermal differentiation such as OTX2, ASCL1, and ZIC1, but were not enriched for PAX motifs (Figure 6L). Conversely, NFR that gained accessibility following BRG1 depletion were most strongly enriched for CTCF/CTCFL motifs and also enriched for KLF/SP1 motifs (Figure 6M). Thus, BRG1 was critical for the maintenance of chromatin accessibility at enhancers likely involved in neuroectoderm differentiation.

### BRG1 establishes the neural progenitor cell chromatin landscape

Performing ATAC-seq at both day 0 and day 3 of NPC differentiation allowed us to examine chromatin accessibility dynamics between transitioning cell types. In control ESCs, 16,723 NFR lost accessibility during early NPC differentiation, and 18,217 NFR gained accessibility. We labelled these NFRs as Day3-lost and Day3-gained NFR respectively (Figure 7A). The same statistical comparison in BRG1-depleted cells revealed a modest decrease in the number of Day3- lost NFR, and a greater decrease in the number of Day3-gained NFR (Figure 7B). This demonstrated that BRG1 was required for chromatin accessibility dynamics during NPC differentiation.

**Figure 7:**
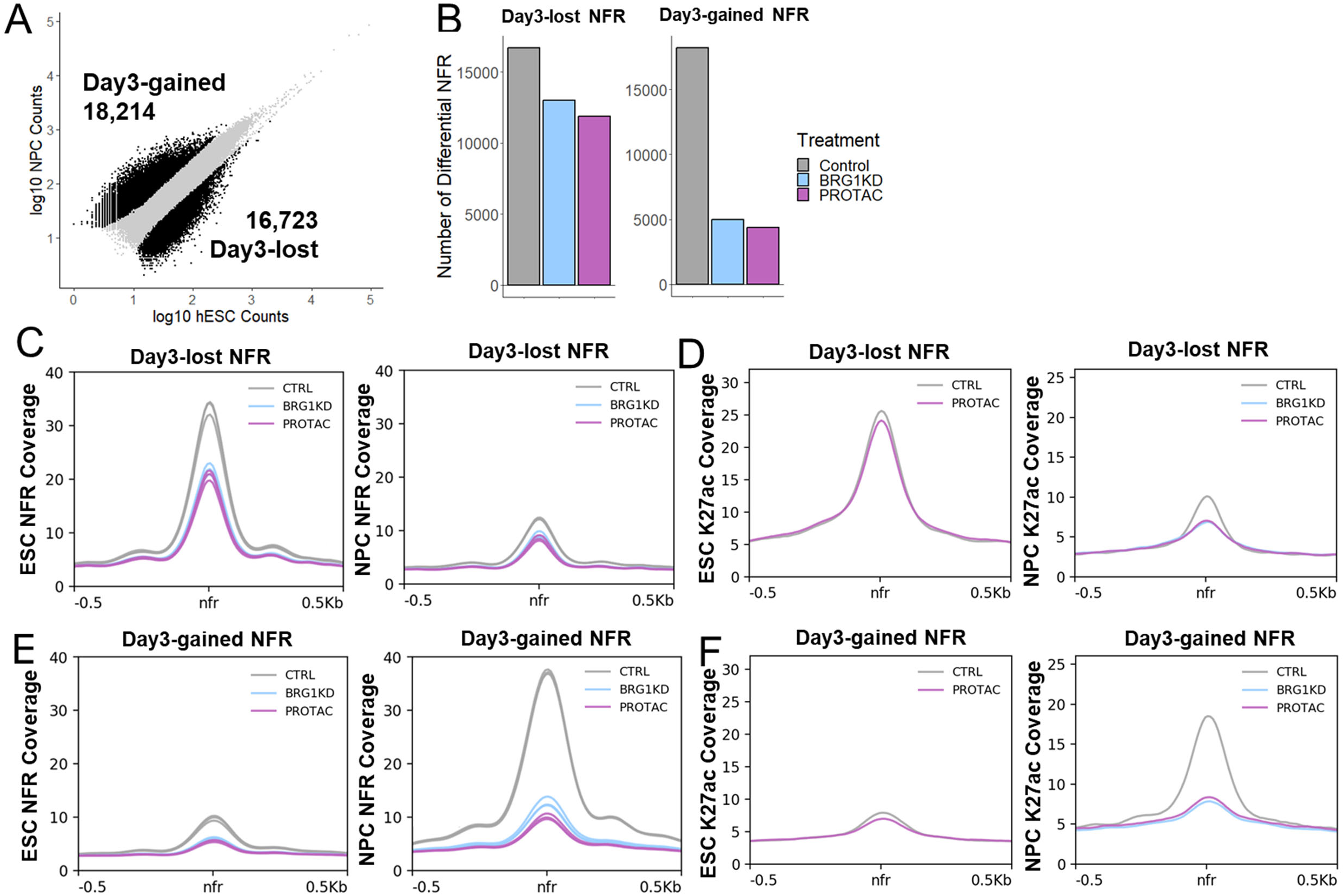
BRG1 establishes the neural progenitor cell chromatin landscape. A) Scatter plot comparing NFR counts between ESC and NPCs over the total set of NFR. Black dots and numbers represent NFR with significantly more or less accessibility in NPCs vs ESCs. B) Bar plots depicting the number of Day3-lost and Day3-gained NFR in each ESC-NPC treatment pair. C) Meta-profiles of ESC NFR and NPC NFR coverage over Day3-lost NFR. D) Meta- profiles of ESC and NPC H3K27ac Cut&Tag coverage over Day3-lost NFR. E) Meta-profiles of ESC NFR and NPC NFR coverage over Day3-gained NFR. F) Meta-profiles of ESC and NPC H3K27ac Cut&Tag coverage over Day3-gained NFR.

Despite lower levels of accessibility at day 3, BRG1 depletion reduced accessibility at Day3-lost NFR in both ESCs and NPCs (Figure 7C). H3K27ac was also decreased at Day3-lost NFR in NPCs and even further decreased after BRG1-depletion (Figure 7D). Thus, BRG1-dependent changes in accessibility and acetylation at Day3-lost NFR were in the same direction as the changes that occurred during differentiation. As such, we reasoned that the overall loss of accessibility at these NFR in ESC resulted in lower fold changes in accessibility during differentiation and thus fewer statistically significant Day3-lost NFR in BRG1-depleted cells. This indicated that BRG1 was not required for the loss of chromatin accessibility at NFR that “closed” during NPC differentiation.

At Day3-gained NFR, the requirement for BRG1 was more obvious. Again, accessibility was reduced at Day3-gained NFR in both ESC and NPCs (Figure 7E). However, the differentiation-associated increase in accessibility at these NFR was markedly reduced in BRG1-depleted cells (Figure 7E). The Day3-gained NFR had low levels of H3K27ac enrichment in ESC and gained H3K27ac enrichment during differentiation (Figure 7F). Conversely, BRG1 depletion blocked the accumulation of H3K27ac signal at Day3-gained NFR (Figure 7F). Thus, BRG1 was required for differentiation-associated increases in both chromatin accessibility and H3K27ac at Day3-gained NFR. Taken together, these experiments demonstrated that BRG1 was required to “open” chromatin and activate neuroectodermal enhancers during NPC differentiation.

## DISCUSSION

The BAF complex is essential throughout many stages of mammalian development and the integrity of the BAF complex is critical for human health^5, 7, 42^. Specific requirements for the BAF complex in neural development have been demonstrated with a variety of genetic disruptions of individual BAF complex subunits^30^. We previously demonstrated that the core subunit BAF47 (SMARCB1) was required in ESC for enhancer regulation and neuroectodermal differentiation^23^. However, depletion of BAF47 did not disrupt the formation of the BAF complex and, as such, the requirement for BAF complex function in neuroectodermal differentiation remained to be elucidated.

Here, we directly address the role of the BAF complex by depleting BRG1, the predominant catalytic subunit in ESC. BRG1 depletion promoted up-regulation of genes enriched for activation of basic processes related to neural development including several transcription factors involved in neuroectodermal differentiation. During directed differentiation to NPCs, depletion of BRG1 disrupted the normal process of lineage specification. While control NPCs acquired a forebrain-like NPC fate, BRG1-depleted NPCs exhibited abnormal lineage trajectories. BRG1-depleted NPCs exhibited gene expression patterns associated with progenitor cells in the dorsal neural tube, were more prone to neuronal differentiation, and precociously gave rise to neural crest cells. These effects were only observed if BRG1 was depleted during the first three days of differentiation. Thus, BRG1 was required during NPC differentiation to prevent NPC dorsalization and inappropriate neural crest specification.

BRG1 depletion disrupted the normal expression patterns of NPC-enriched transcription factors such as SOX2, PAX6, POU3F1, SOX21, and LHX5 and resulted in the formation of a heterogeneous NPC population with multiple distinct cell fates. BRG1-depleted NPCs predominantly adopted a dorsalized cell fate largely marked by expression of transcription factors such as PAX3 and ZIC1. A neural crest cell population marked by expression of the transcription factors SOX10, FOXD3, ETS1, and TFAP2A/B/C also arose from BRG1-depleted NPCs. BRG1- depleted NPCs were also more prone to neuronal differentiation and formed a differential neuronal population with enhanced expression of the transcription factor ASCL1. Expression of these lineage-specific transcription factors likely drove the emergence of these cell populations. However, the NFRs lost after BRG1 depletion were also enriched for motifs of many of these factors. Furthermore, many of these transcription factors have been shown to recruit BRG1 and require BRG1 to function in a variety of contexts^32, 33, 43, 44^. Therefore, the role of these transcription factors in directing cell fate decisions following BRG1 depletion remains unclear. Directly interrogating their genomic distribution and requirement in the distinct NPC cell clusters could reveal how depletion of BRG1 alters their roles in regulating lineage-specific transcriptional programs.

In ESC and day 3 NPCs, BRG1 was essential to maintain accessible chromatin at thousands of nucleosome-free regions. The NFR with BRG1-dependent accessibility tended to be distal from annotated TSSs and were marked by H3K27ac, indicating that they were likely active enhancers. Depletion of BRG1 reduced both accessibility and H3K27ac at BRG1-dependent sites and these sites were strongly enriched for SOX motifs and motifs of other neuroectodermal transcription factors. Thus, it appeared that BRG1 was necessary to maintain accessibility and histone acetylation at enhancers involved in NPC differentiation. Indeed, BRG1 depletion greatly reduced the number of genomic regions that gained accessibility during the first three days of NPC differentiation. More than 10,000 sites that gained accessibility and histone acetylation during differentiation were dependent on BRG1. Together, these findings demonstrated that BRG1 was required to open and activate chromatin at enhancers essential for normal NPC specification.

The enrichment of SOX family transcription factor motifs within BRG1-dependent NFR suggested that BRG1 was vital for a SOX-driven transcriptional program during NPC differentiation. SOX transcription factors directly interact with BAF complexes in a variety of contexts including NPCs and other neural lineages. Interestingly, SOX factors can function as pioneer factors and bind to nucleosomal DNA to promote DNA accessibility for other factors^45^. Other pioneer factors such as OCT4 and PU.1 require BRG1 to generate accessible chromatin^46, 47^. As such, we propose a salient hypothesis for BRG1 function during NPC specification: activation of NPC transcriptional programs is dependent on BRG1-mediated chromatin remodeling directed by SOX transcription factors.

The different effects of BRG1 and BAF47 depletion on NPC specification also provide insight into the regulation of BAF complex function during development. We and others have shown that BAF47 depletion impairs neural differentiation from human ESC^23, 48^. Despite varied reports on the role of BAF47 in the composition and integrity of the BAF complex, BAF47 depletion or deletion is necessary for BAF complex interaction with enhancers, regulation of enhancer activity, eviction of PRCs from chromatin, and activation of bivalent promoters^49–54^. Thus, depletion of BAF47 impairs or alters BRG1 and BAF complex function. The effects of this impairment or alteration on neuroectodermal differentiation are distinct from the effects of BRG1 loss or depletion. These findings serve to demonstrate the intricacies and difficulties of targeting BAF complex activity. As such, consideration must be given to context-dependent distinctions between different modalities for targeting BRG1 and the BAF complex for therapeutic purposes.

BRG1 and BAF subunit mutations have been identified in multiple aggressive pediatric brain tumors. Atypical teratoid/rhabdoid tumors (ATRTs) are commonly associated with mutations in BAF47, but a small and distinct subclass of ATRTs have bi-allelic mutations in BRG1 ^18, 20^. Tumors of this ATRT-SMARCA4 subgroup have higher incidence of germline mutations, earlier age-of-incidence, and worse prognoses than the other subgroups with BAF47 mutations ^18, 20^. BRG1 mutations are also frequently found in the WNT-group and Group-III subgroups of medulloblastomas, with Group-III being the subgroup with the earliest age-of-incidence and worst prognoses^55, 56^. Both tumor types are thought to arise from neural progenitors in the midbrain, hindbrain, or neural crest. Intriguingly, only BRG1KD NPCs were classified with these cell-type annotations when modeled on transcriptional profiles from first-trimester fetal human brains. Thus, we propose that loss of BRG1 function during the earliest stages of differentiation causes aberrant patterning of NPCs and yields a greater proportion of cells with dorsal/caudal fates. This aberrant patterning may result in these NPCs being exposed to inappropriate signaling environments with the potential to promote tumorigenesis. Intriguingly, BAF47 depletion at later stages of neural differentiation leads to failures in neuronal maturation and produces NPCs with a transcriptional profile similar to ATRTs^48^. Tracking BRG1-depleted NPCs through later stages of differentiation to determine their ultimate cell fates could provide greater insight into the pathogenicity of BRG1 mutations during embryogenesis.

## Limitations of this study

In this study, we focus on the initial stages of NPC differentiation using the dual-SMAD inhibition protocol. It would be interesting to examine the effects of BRG1 depletion on other neuroectodermal differentiation protocols. It would also be interesting to use more directed protocols for specific neuroectodermal lineages (eg specific types of neurons, glia, or neural crest cells). Additionally, while we propose potential mechanisms and highlight several transcription factors that may direct BRG1 activity, we do not extend our study to test these hypotheses. Testing the effects of BRG1 depletion on candidate transcription factor binding profiles could provide mechanistic evidence for the molecular activity of the BAF complex in cell fate decisions.

## Author Contributions

Conceptualization, JAH, GWM, and TKA; Methodology, JAH, GWM, and LFL; Investigation, JAH, GWM, and LFL; Validation, JAH, GWM, and IG; Formal Analysis, JAH and IG; Visualization, JAH; Writing – Original Draft, JAH and GWM; Writing – Reviewing & Editing, JAH, GWM, and TKA; Funding Acquisition, TKA

## STAR★Methods

### Key resources table

| REAGENT or RESOURCE | SOURCE | IDENTIFIER |
| --- | --- | --- |
| **Antibodies** |  |  |
| BRG1 | Santa Cruz Biotechnologies | sc-17796, RRID:AB_626762 |
| BRG1 | Novus Biologicals | NB100-2594, RRID:AB_2191852 |
| Actin | Santa Cruz Biotechnologies | sc-10731, RRID:AB_2223515 |
| SOX2 | Millipore | AB5603, RRID:AB_2286686 |
| PAX3 | DSHB | Pax3, RRID:AB_2315061 |
| LHX5 | DSHB | PCRP-LHX5-1B7, RRID:AB_2722232 |
| BAF60A | BD Biosciences | 611728, RRID:AB_2192143 |
| Beta-tubulin | DSHB | E7, RRID:AB_528499 |
| OCT4-conjugated | BD Biosciences | 560329, RRID:AB_1645318 |
| SOX2-conjugated | BD Biosciences | 562139, RRID:AB_10897844;<br>561593, RRID:AB_10894382 |
| PAX6-conjugated | BD Biosciences | 562388, RRID:AB_11153319 |
| NGFR-conjugated | BD Biosciences | 562562, RRID:AB_2737657 |
| H3K27ac | Abcam | ab4729, RRID:AB_2118291 |
| Anti-Rabbit CUT&Tag Secondary | Epicyphe | 13-0047 |
| IRDye 800CW Donkey anti-Rabbit IgG | LI-COR Biosciences | 926-32213, RRID:AB_621848 |
| IRDye 680RD Donkey anti-Rabbit IgG | LI-COR Biosciences | 926-68073, RRID:AB_10954442 |
| IRDye 800CW Donkey anti-Mouse IgG | LI-COR Biosciences | 926-32212, RRID:AB_621847 |
| IRDye 680RD Donkey anti-Mouse IgG | LI-COR Biosciences | 926-68072, RRID:AB_10953628 |
| **Chemicals, peptides, and recombinant proteins** |  |  |
| Formaldehyde Solution, 16% | ThermoFisher Scientific | 28906 |
| TeSR-E8 Basal Medium | STEMCELL Technologies | 05990 |
| hESC Matrigel | Corning | 354277 |
| ReLeSR reagent | STEMCELL Technologies | 05872 |
| Y-27632 | STEMCELL Technologies | 72304 |
| TrypLE Express reagent | Gibco | 12604013 |
| STEMdiff™ SMADi Neural Induction Kit | STEMCELL Technologies | 08581 |
| ACBI1 PROTAC | MedChemExpress | HY-128359 |
| BRM/BRG1 ATP Inhibitor-1 | MedChemExpress | HY-119374 |
| Doxycycline | Alfa Aesar | J60579 |
| HALT Protease Inhibitors | ThermoFisher Scientific | 78430 |
| TWEEN 20 | Millipore Sigma | P7949 |
| CUTANA pAG-Tn5 | Epicyphe | 15-1017 |
| NEBNext High-Fidelity 2X PCR Master Mix | New England BioLabs | M0541L |
| **Critical commercial assays** |  |  |
| Zombie NIR Fixable Viability Kit | BioLegend | 423105 |
| eBioscience Foxp3 / Transcription Factor Staining Buffer Set | ThermoFisher Scientific | 00-5523-00 |
| AMPure XP beads | Beckman Coulter | A63881 |
| SuperScript III First-strand kit | ThermoFisher Scientific | 18080-051 |
| ssoAdvanced Universal SYBR Green Supermix | Bio-Rad | 172-5274 |
| RNeasy Mini Kit | Qiagen | 74104 |
| QiaQuick PCR Purification Kit | Qiagen | 28104 |
| Dounce Homogenizer | Duran Wheaton Kimble | 357542 |
| Protein A/G Dynabeads | ThermoFisher Scientific | 88802 |
| Bio-Rad Protein Assay Kit II | BIO-RAD | 5000002 |
| Total RNA Purification Plus Kit | Norgen Biotech | 48300 |
| Tagment DNA TDE1 Enzyme Kit | Illumina | 20034197 |
| Chromium Single Cell 3' Library & Gel Bead Kit v3 | 10X Genomics | 1000268 |
| BioMag Plus Concavalin A Beads | Bangs Laboratories Inc. | BP531 |
| **Experimental models: cell lines** |  |  |
| H1 human embryonic stem cells | WiCell | WA01 |
| **Software and algorithms** |  |  |
| Cutadapt 1.9.1 | Martin et al.<sup>57</sup> | RRID:SCR_011841 |
| Bowtie2 2.0.0-beta7 | Langmead and Salzberg<sup>58</sup> | RRID:SCR_016368 |
| Samtools 0.1.20 | Li et al.<sup>59</sup> | RRID:SCR_002105 |
| Picard-tools 1.110(1752) | https://broadinstitute.github.io/picard/ | RRID:SCR_006525 |
| MACS2 2.1.1.20160309 | Zhang et al.<sup>60</sup> | RRID:SCR_013291 |
| Bedtools 2.24.0 | Quinlan and Hall<sup>61</sup> | RRID:SCR_006646 |
| Deeptools 2.5.3 | Ramirez et al.<sup>62</sup> | RRID:SCR_016366 |
| ggplot2 3.4.0 | https://ggplot2.tidyverse.org | RRID:SCR_021139 |
| FlowJo | BD Biosciences | RRID:SCR_008520 |
| Salmon 1.3.0 | Patro et al.<sup>63</sup> | RRID:SCR_017036 |
| Tximport 1.14.2 | Soneson et al.<sup>64</sup> | RRID:SCR_016752 |
| Limma-voom 3.42.2 | Ritchie et al.<sup>65</sup> | RRID:SCR_010943 |
| featureCounts 1.5.0-p2 | Liao et al.<sup>66</sup> | RRID:SCR_012919 |
| DESeq2 1.38.3 | Love et al.<sup>67</sup> | RRID:SCR_015687 |
| CellRanger 3.0.1 | 10X Genomics |  |
| Seurat 4.3.0 | Hao et al.<sup>68</sup> | RRID:SCR_016341 |
| R 3.6.1, 4.2.1, 4.2.2, 4.2.3, and 4.3.1 |  |  |

## Resource availability

### Lead contact

The lead contact is Jackson A Hoffman. Please address correspondence to.

### Materials availability

Materials generated for or used within this study are available upon correspondence.

### Experimental model and subject details

H1 human embryonic stem cells.

## Method details

### ESC cell culture and generation of BRG1KD ESC cell line

ESCs were cultured feeder-free in TeSR-E8 Basal Medium, on Matrigel coated tissue culture vessels, and incubated in a 5% CO2 atmosphere and at 37°C. ReLeSR reagent was used to collect ESC aggregates during general passaging. Sub-cloning of the shRNA against SMARCA4 was performed by digestion of the pINDUCER10 back-bone vector with Xho I and Mlu I, following standard procedures as previously described^23, 69^. Lentiviruses carrying the respective shRNAs were produced at the NIEHS Viral Vector Core Laboratory according to a previously established protocol^70^. H1 cells were infected at MOI8 and selected using 1 mg/ml puromycin for 24 hr. After an 18hr 1ug/ml doxycycline treatment, RFP+HIGH cells were used to establish the shBRG1 cell line and used for all subsequent experiments. Doxycycline treatment of ESC cultures for induction of shRNA and RFP expression was performed using 1ug/ml doxycycline in TeSR-E8 media for 3 days, unless otherwise stated. ACBI1 PROTAC treatment for targeted degradation of BRG1 was performed using 0.5uM PROTAC in TeSR-E8 media for 24 hours, unless otherwise stated.

### Monolayer neural induction protocol and hNPC culture

ESCs were cultured as described and collected as single cell suspensions using the TrypLE Express reagent. Cells were then cultured according to the STEMCELL Technologies instructions for the STEMDiff Dual Neural Induction System for the generation of neural progenitor cells (NPC) using dual SMAD inhibition. Briefly, ESCs were plated as single cells at a density of 2.0E05cells/cm2 in STEMDiff Neural Induction Media with SMADi, and 10uM Y- 27632, on Matrigel coated plates, and incubated in a 5% CO2 atmosphere at 37°C. Media was changed daily with STEMDiff Neural Induction Media with SMADi. In experiments where shRNA expression was induced using doxycycline or BRG1 protein degradation was induced using the PROTAC molecule, doxycycline or PROTAC was included in the media throughout the experiment unless otherwise indicated. At specific experimental time points during differentiation, NPCs were collected using Accutase and further processed depending on the downstream application.

### Quantitative PCR

RNA was isolated from cells using RNeasy Plus kit purification with gDNA eliminator columns. cDNA was generated from total RNA using SuperScript III and either oligo(dT) or random hexamers. qPCR was run using ssoAdvanced Universal SYBR Green Supermix and gene/transcript specific primers. All experiments were performed with 3 or more biological replicates. qPCR data was normalized to control genes.

### Western blotting

Cells were lysed in TINE300 buffer (100mM Tris-HCL pH8.0, 1% IGEPAL CA-630, 300mM NaCl, 1mM EDTA, 0.25mM EGTA) containing 1:100 protease inhibitor cocktail and 1mM PMSF. Cells were allowed to lyse for 20min on ice followed by 30s of sonication. Lysate total protein concentrations were estimated by Bio-Rad protein assay. 20-40ug of total protein was applied per lane of acrylamide tris-glycine gels run at 125V. Proteins were then transferred from gel to nitrocellulose membranes. Blots were then blocked for 30min at room temperature in blocking solution (1XTBS with 2% milk), followed by an overnight incubation in blocking solution plus primary antibodies. Blots were washed with wash buffer (1XTBS with 1% Tween-20), followed by incubation in blocking buffer with species appropriate secondary antibodies. Blot imaging was performed using the LI-COR Odyssey imager.

### Flow cytometry

H1 ESCs and NPCs single cells were collected using dissociation protocols described above, washed with 1XPBS and stained with fixable viability dye. Cells were washed with FACS Buffer (1XPBS, 2%BSA, 1mM EDTA, 0.1%NaAzide), followed by incubation in FACS buffer with fluor-conjugated surface antibodies for 2 hours at room temperature and with occasional mixing.

Cells were washed with FACS Buffer and incubated in fixation and permeabilization buffer at 4°C overnight. Fixed cells were washed with permeabilization wash buffer followed by incubation in permeabilization wash buffer with intracellular target antibodies. Cells were washed with FACS Buffer prior to analysis using a Becton Dickinson LSRFortessa flow cytometer. Flow cytometry data was processed using FlowJo v10. Debris, cell clusters, and permeability dye positive cells were excluded from downstream analysis. Single marker quantification was performed using histogram gating as compared to known positive populations and isotype controls (Figure S3A). Graphed data presented has mean values of 2 or more independent experiments with 3 or more biological replicates each.

### RNA-seq

RNAseq experiments were performed on 3 biological replicates per sample. RNA was isolated from ESCs using the total RNA purification kit followed by ribosome RNA removal, library preparation, and sequencing by Expression Analysis, an IQVIA company. Transcript abundance was quantified using Salmon^63^ with an index containing Gencode version v32lift37 comprehensive transcripts^71^, human Class I and II enhancers^35^, and the hg19 human genome as decoy. Data were imported into R-3.6.1 for analysis, summarized to gene level with tximport^64^, then statistical contrasts were performed using limma-voom^65^ moderated t-tests.

### ATAC-seq

Single cell suspensions of H1 ESCs or day 3 NPCs were generated as described above. Tagmentation was performed on 50,000 cells using Tn5 enzyme in tagmentation buffer on 3 biological replicates per condition. Isolation of libraries and amplification was performed as described^72^. Paired-end reads were trimmed with cutadapt^57^, aligned to hg19 with bowtie2^58^, and deduplicated with picardtools. Nucleosome free reads were defined with bamtools by keeping reads with insert sizes less than 101 bases. NFR were identified by calling peaks with macs2^60^ with mfold limits of 10 and 200 and FDR cutoff of 0.001. NFR were merged with bedtools^61^ and filtered to only keep NFR that were called in more than one replicate of at least one treatment group. Nucleosome free reads within NFR were quantified with FeatureCounts^66^ and differential analysis was performed with DESeq2^67^ with fold change > 2 and adjusted p-value < 0.01 cutoffs.

### Single-cell RNA-seq

Single cell suspensions were prepared for 2 biological replicates per cell type at concentration of 0.8-1.2x10^6^ cells/ml and with cell viability of 80% or above. Briefly, approximately 5000 cells were used together with the Chromium Single Cell 3’ Library & Gel Bead Kit v3.1 onto the 10x single cell chromium chip to generate a single cell emulsion in Chromium Controller. Reverse transcription of mRNA and cDNA amplification were carried out following the manufacture’s instruction. The amplified cDNA was further fragmented to construct NGS libraries. The libraries were then sequenced by the NIEHS Epigenomics and DNA Sequencing Core Laboratory with the parameters recommended in the manufacturer’s instruction. Fastq files were processed with CellRanger v3.0.1 using GRCm38 genome and Gencode annotation provide by 10X to generate the initial cell by gene matrix for each sample. Matrices were read into Seurat v4^68^ using R-4.2.1 and merged into a single Seurat object. Cells with greater than 10% mitochondrial reads or less than 4000 UMI were removed from the object. UMAP dimensional reduction was performed using the top 15 principal components and clusters were called with resolution = 1.5. From this object, the raw counts matrix was exported for cell annotation using the online tool at celltypist.org. Data were visualized with built-in Seurat tools or with ggplot2.

### Cut&Tag

Cut&Tag was largely performed as per Epicypher and Henikoff lab protocols. Briefly, nuclei were isolated, fixed for 2 minutes with 0.1% Formaldehyde at room temperature, and cryopreserved prior to all Cut&Tag experiments. For each experiment, nuclei were thawed and 100,000 nuclei were used per reaction. Nuclei were bound to activated Concanavalin A beads and 0.5ug of primary antibody was bound overnight at 4C. Secondary antibody was bound for 30 minutes at room temperature, pAG-Tn5 was bound for 1 hour at room temperature, and the tagmentation reaction was incubated at 37C for 1 hour. Libraries were amplified with NEBNext high-fidelity PCR mix, cleaned-up with AMPure XP beads, and sequenced by the NIEHS Epigenomics and DNA Sequencing Core Laboratory. Paired-end reads were trimmed with cutadapt^57^, aligned to hg19 with bowtie2^58^, and deduplicated with picardtools. Coverage files and meta-profiles were generated with Deeptools^62^.

## Acknowledgements

We are grateful to the members of the Archer lab, the NIEHS Epigenetics and Stem Cell Biology Laboratory, the NIEHS Integrative Bioinformatics Group, and Dr. David Fargo for their ongoing support, advice, and constructive criticism. We also thank Greg Solomon, Jason Malphurs, Nicole Reeves, and Xin Xu of the NIEHS Epigenomics Core Laboratory for their next generation sequencing expertise. Finally, we thank Dr. Paul Wade, Dr. Joe Rodriguez, and Dr. Carl Bortner for critical review of the manuscript and data. This work was supported by funding from the National Institutes of Environmental Health Sciences (Z01-ES071006-23).

## Declaration of interests

The authors declare no competing interests.

## Data and code availability

• All sequencing data generated for this study will be deposited at GEO

• This paper does not report original code.

• Any additional information required to reanalyze the data reported in this paper is available from the lead contact upon request.

## SUPPLEMENTAL FIGURE LEGENDS

**Figure S1:**
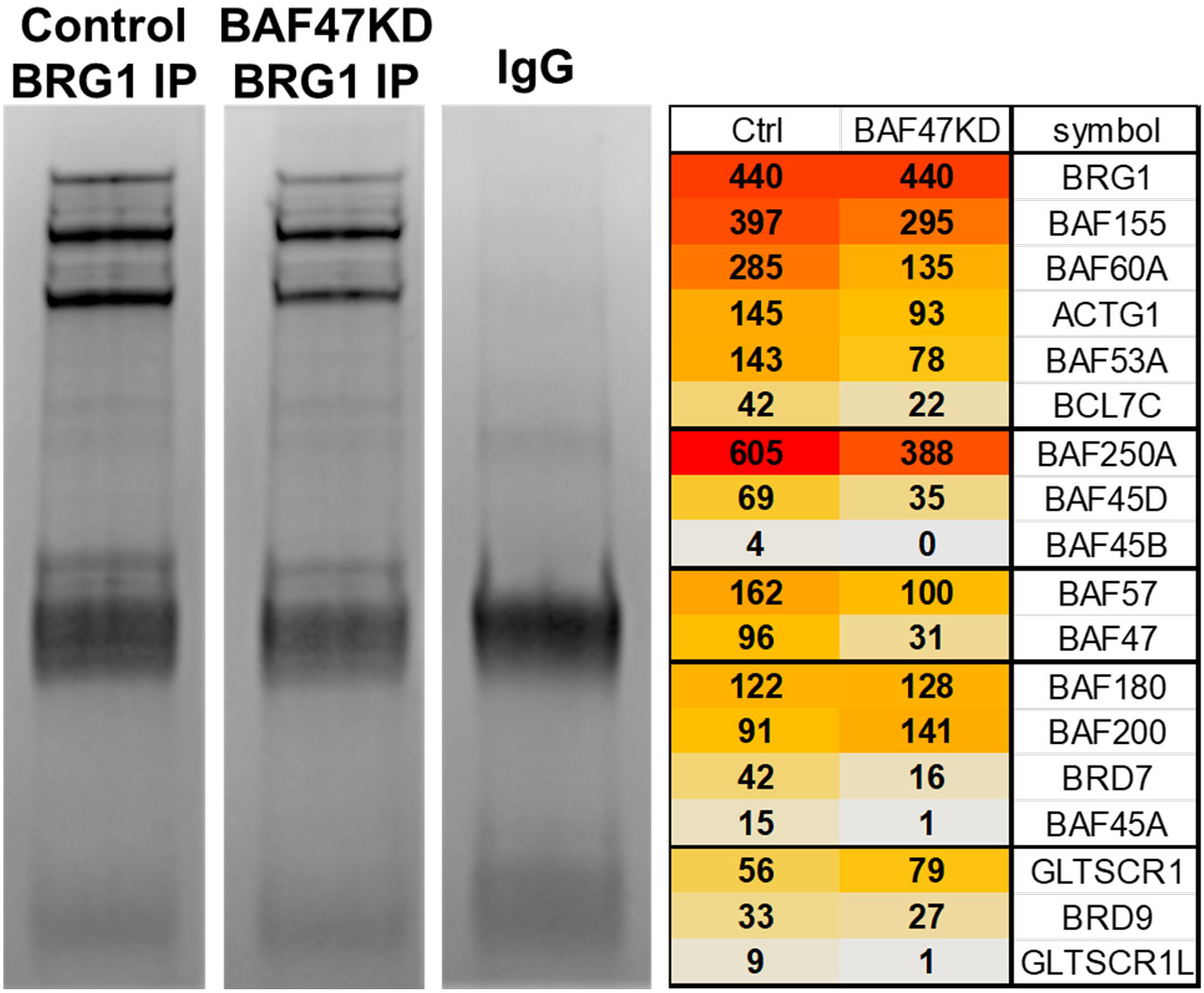
BAF47-independent formation of BAF complexes. Coomassie stained images of BRG1 or IgG immunoprecipitations from control or BAF47KD ESC. Table shows relative number of spectra detected for BAF complex subunits. Spectra counts normalized to number of BRG1 spectra.

**Figure S2:**
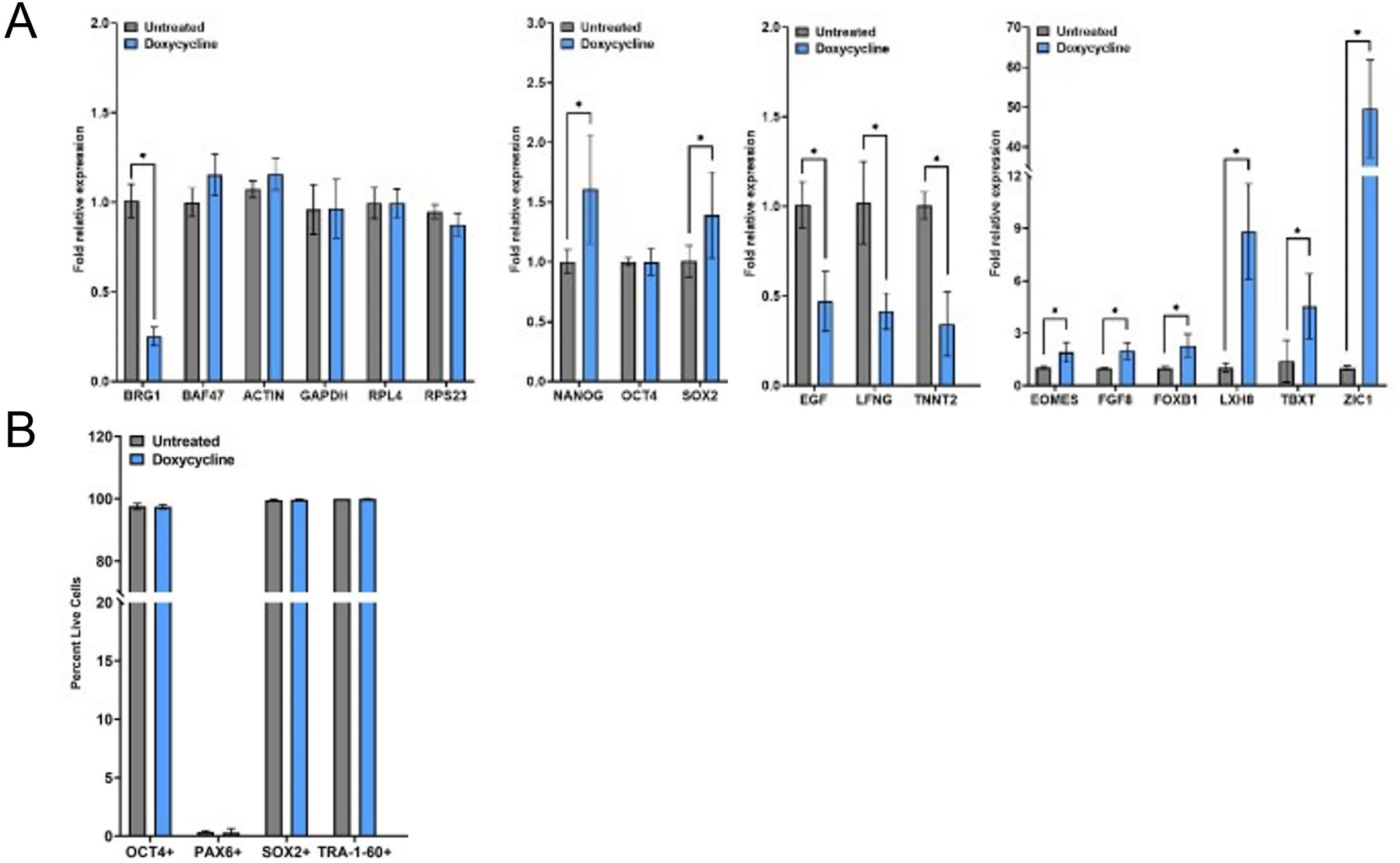
Altered gene expression in BRG1KD ESC. A) RT-PCR for BAF subunits, pluripotency genes, down-regulated RNA-seq DEGs, and up-regulated RNA-seq DEGs. B) Percent of control or BRG1KD ESC positive for the listed markers.

**Figure S3:**
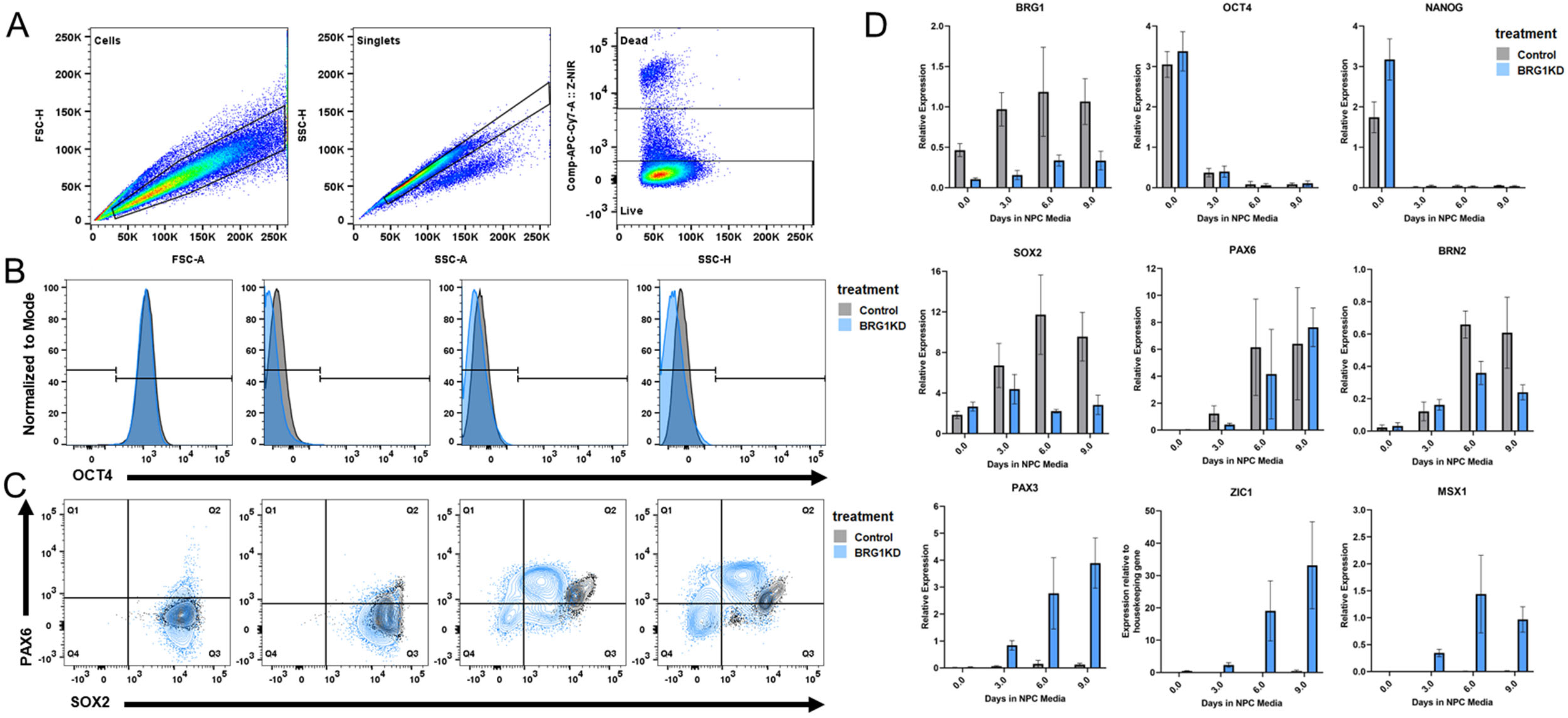
Altered protein and gene expression in BRG1KD NPCs. A) FACS gating parameters. B) FACS histogram of OCT4 protein detection at collection timepoints. Log10 fluorescent signal depicted for control cells in gray and BRG1KD cells in light blue. Data represents the total of all biological replicates (n >= 3). C) FACS contour plot of SOX2 and PAX6 protein detection at collection timepoints. Contour lines represent 2% intervals of cell density, control cells in gray and BRG1KD cells in light blue. Data represents the total of all biological replicates (n >= 3). D) RT-PCR for BRG1, pluripotency markers, and neural transcription factors. Bars depict mean expression level relative to control genes; error bars represent standard deviation of biological triplicates.

**Figure S4:**
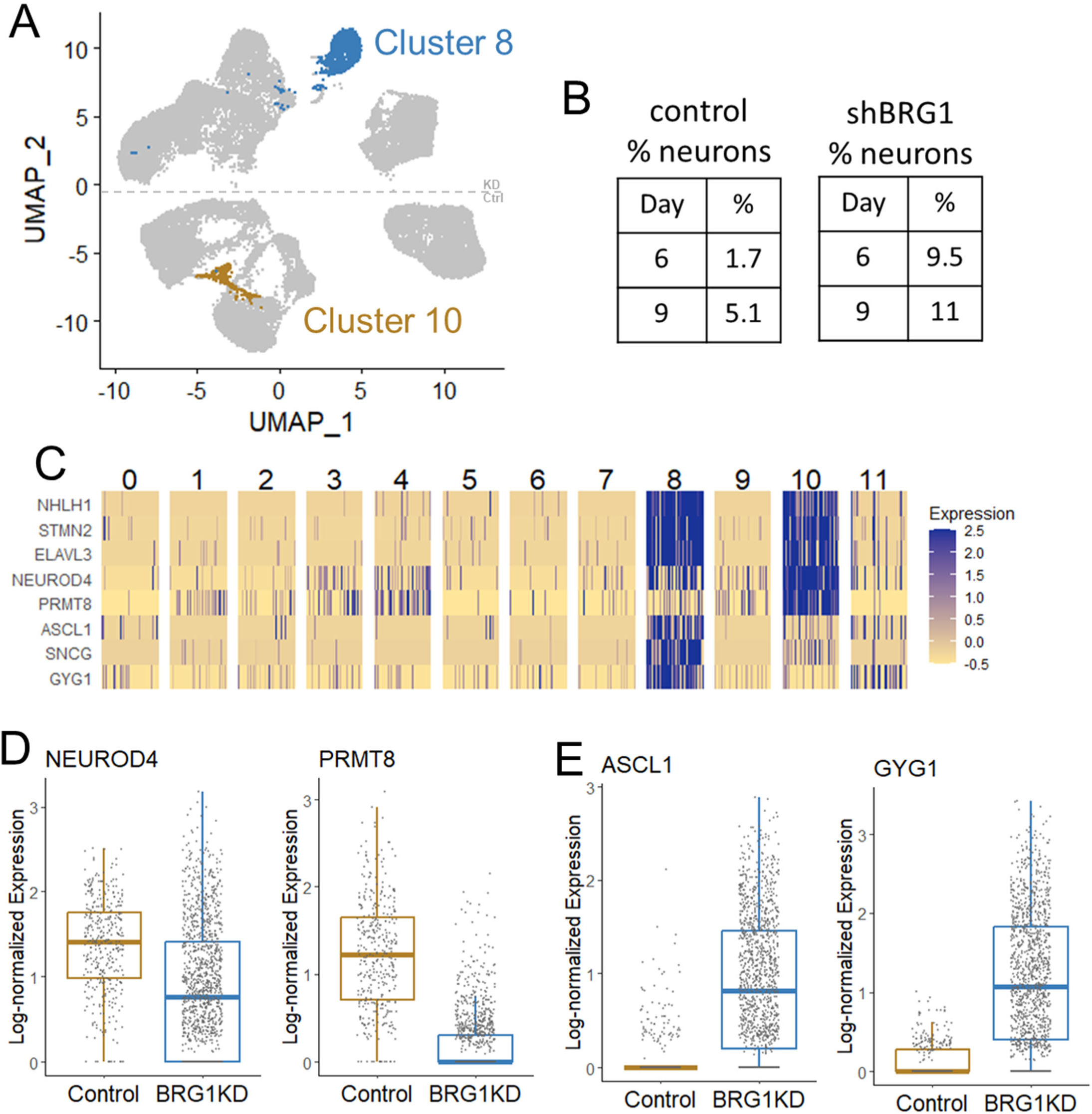
Altered neuronal specification in BRG1-depleted NPCs. A) UMAP plot highlighting clusters 8 and 10. B) Percentages of control and BRG1KD cells in clusters 10 and 8, respectively. C) Heatmap of log-scaled expression of neuronal marker genes in randomly down-sampled Seurat clusters (50 cells per cluster). D,E) Box and jitter plots of neuronal marker genes differentially expressed between clusters 8 and 10.

**Figure S5:**
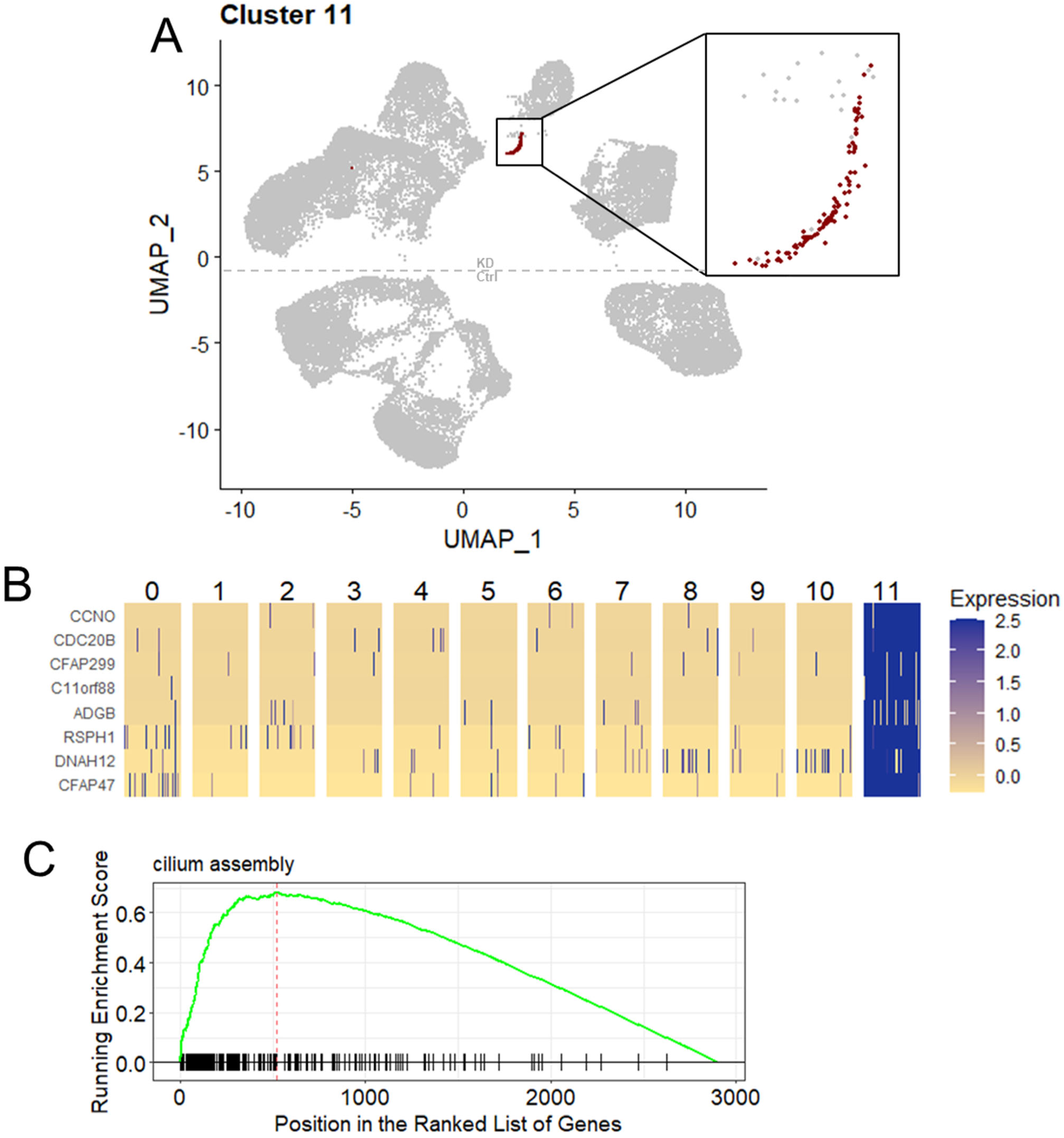
Precocious ciliated cell specification from BRG1-depleted NPCs. A) UMAP plot highlighting cluster 11. B) Heatmap of log-scaled expression of ciliated cell marker genes in randomly down-sampled Seurat clusters (50 cells per cluster). C) GSEA plot of running enrichment score for “cilium assembly” gene set within assigned marker genes of cluster 11.

**Figure S6:**
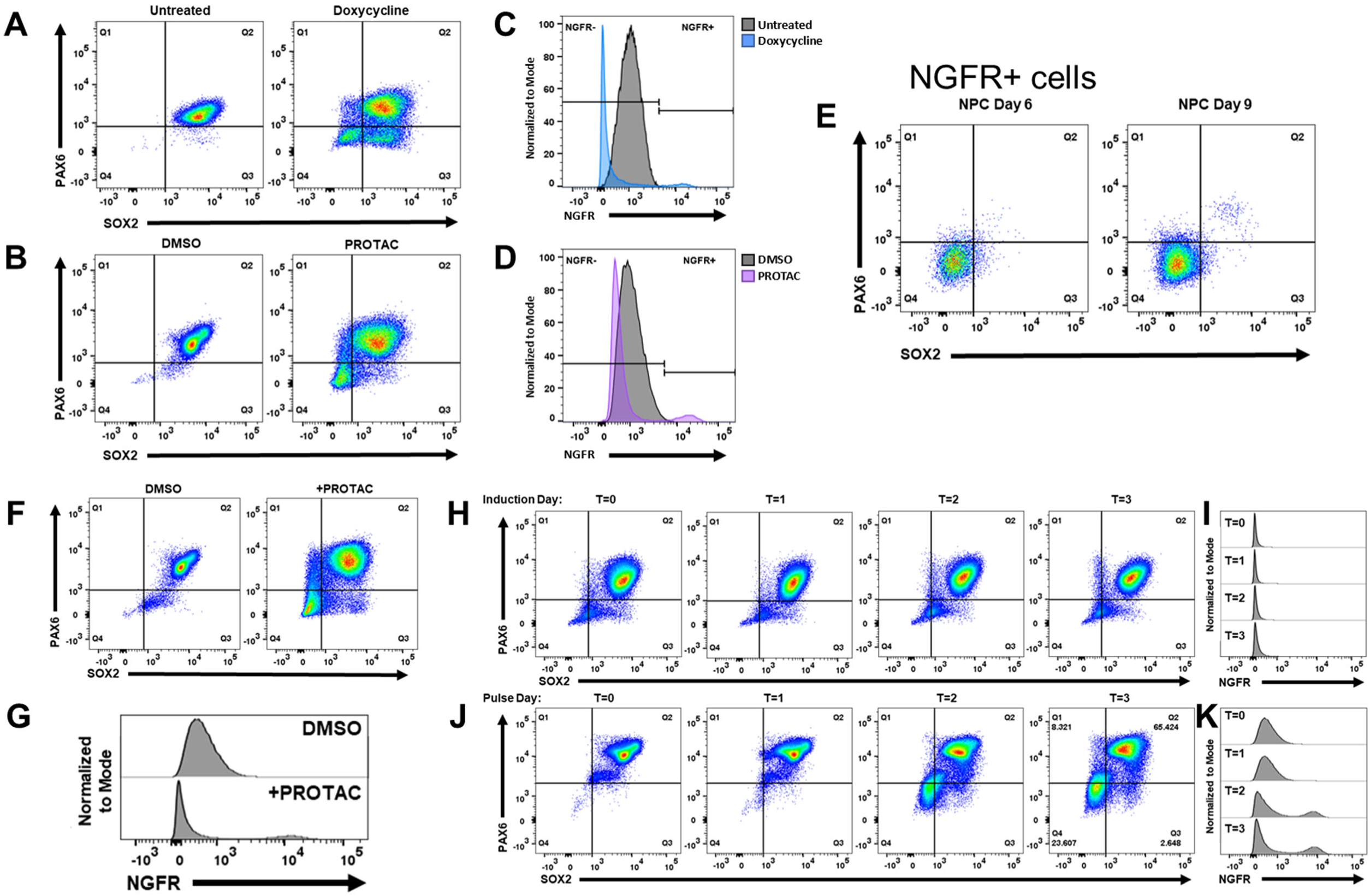
FACS scatter plots corresponding to. Figure 5. A,B,F,H, and J) Cell scatter plots depicting levels of SOX2 and PAX6 protein expression. Color indicates density of points; blue = low, red = high. C,D) FACS histograms depicting the density of cells expressing NGFR protein. F) Cell scatter plots depicting levels of SOX2 and PAX6 protein expression in NGFR+ BRG1KD NPCs at days 6 and 9. Color indicates density of points; blue = low, red = high. G, I, and K) FACS histograms of NGFR protein expression.

## Notes

### Competing Interest Statement

The authors have declared no competing interest.

